# Cholesterol in the cargo membrane reduces kinesin-1 binding to microtubules in the presence of tau

**DOI:** 10.1101/2022.03.01.482581

**Authors:** Qiaochu Li, James T. Ferrare, Jonathan Silver, John O. Wilson, Luis Arteaga-Castaneda, Weihong Qiu, Michael Vershinin, Stephen J. King, Keir C. Neuman, Jing Xu

## Abstract

Intracellular cargos are often membrane-bound and transported by microtubule-based motors in the presence of microtubule-associated proteins (MAPs). Whereas increasing evidence reveals how MAPs impact the interactions between motors and microtubules, critical questions remain about the impact of the cargo membrane on transport. Here we combined *in vitro* optical trapping with theoretical approaches to determine the effect of a lipid cargo membrane on kinesin-based transport in the presence of MAP tau. Attaching kinesins to a fluid lipid membrane decreases the inhibitory effect of tau in comparison to membrane-free cargos. Adding cholesterol, which reduces kinesin diffusion in the cargo membrane, amplifies the inhibitory effect of tau on kinesin in a dosage-dependent manner. Our findings can be understood in a framework in which kinesin diffusion in the cargo membrane counteracts binding-site occlusion by tau. Our study establishes a direct link between the physical properties of cargo membrane and MAP-based regulation of kinesin-1.

## Introduction

Kinesin-1 is a major microtubule-based motor protein responsible for long-range transport in eukaryotic cells (1, 2). Underscoring the critical importance of kinesin in cell function and survival, mutations in the neuronal kinesin heavy chain isoform 5A (Kif5A) cause neurodegenerative diseases including hereditary spastic paraplegias (3), Charcot-Marie-Tooth disease type 2 (4), and amyotrophic lateral sclerosis (5). Kinesin transports cargos by processively stepping along microtubules (6, 7); this processive motion is initiated when the motor binds the microtubule and is terminated when it unbinds from the microtubule. Tuning of the binding and unbinding rates of kinesin, for example via the presence of microtubule-associated proteins (MAPs) (8-10), has emerged as a key mechanism of kinesin regulation. In particular, the MAP tau occludes kinesin binding sites on the microtubule (10-12), inhibiting binding and promoting unbinding of kinesin *in vitro* (13-15). However, *in vivo*, tau is essential for microtubule assembly and stabilization (16). How kinesin maintains transport in the presence of tau, particularly in environments such as healthy neurons where tau is enriched, remains to be elucidated.

In cells, kinesins are often attached to their cargos via a fluid lipid membrane (17-19). This cargo-enclosing membrane is absent from most *in vitro* investigations, including the pioneering studies that established the inhibitory effects of tau on kinesin-based transport (13-15). The lipid membrane enclosing cargos *in vivo* is “fluid” in that its constituents, including motor proteins such as kinesin, are mobile and can diffuse on the cargo surface. As membrane fluidity increases, each motor protein can more quickly diffuse to open binding sites on the microtubule via diffusive search. Alterations in membrane cholesterol or other fluidity-restricting lipids are increasingly linked to aging and neurological diseases (20, 21), in which dysfunctions in motor-based transport have emerged as a common early hallmark (22). These *in vivo* observations suggest that diffusion of motors in the cargo membrane may contribute to transport regulation. However, recent investigations found no significant effects of a lipid membrane (23, 24) on the binding or unbinding rates of kinesins to microtubules in the absence of MAPs.

Here we employed *in vitro* optical trapping to determine how a lipid cargo membrane affects kinesin-based transport in the presence of the MAP tau. We hypothesized that diffusion of motors in the cargo membrane counteracts binding-site occlusion by tau, thereby enhancing kinesin binding to microtubules in the presence of tau. To test this hypothesis, we generated membrane-enclosed cargos by enclosing silica microspheres with a fluid lipid bilayer (23, 25, 26). We then added cholesterol to the bilayer to reduce membrane fluidity (24, 27), which in turn reduced kinesin diffusion in the membrane (24, 27). We next developed a theoretical model to provide mechanistic insights into our *in vitro* findings. Together, our experimental and theoretical results support our hypothesis and establish a direct link between the physical properties of cargo membrane and MAP-based regulation of kinesin-1.

## Results

### A fluid lipid membrane reduces the inhibitory effect of tau on kinesin-1

We first tested our hypothesis by comparing the transport of membrane-enclosed versus membrane-free cargos. We used a cholesterol-free, dipalmitoylphosphatidylcholine (DOPC) lipid bilayer to model the fluid membrane of intracellular cargos (26, 28, 29), and we included biotinylated or nickel-loaded chelator lipids in the bilayer to mediate kinesin attachment (Table 1). We employed human tau 23 (htau23, or 3RS tau), an isoform of tau that strongly inhibits kinesin *in vitro* (13-15), and we characterized cargo transport by ∼1-2 kinesins as observed for intracellular vesicles (17-19). To quantify transport, we used a custom-built optical trap to position individual kinesin-cargo complexes near a microtubule for up to 30 s (Fig. 1*A, Inset*); this 30-s interaction time is sufficient to observe kinesin binding to microtubules in the absence of tau (14, 30, 31). We scored the fraction of cargos that bound to, and moved along, the microtubule (“motile fraction”). For cargos that were motile, we turned off the optical trap and measured the distance that the cargo traveled before unbinding from the microtubule (“run length”). Both motile fraction and run length impact the flux of cargos transported *in vivo*; they are also appropriate for probing the hypothesized change in motor binding to the microtubule. Specifically, for processive motors such as kinesin-1, motile fraction reflects the binding rate of the motor (14, 30, 31), whereas run length is more nuanced and can depend on both the unbinding and the binding rates of the motor (32, 33). Because the inhibitory effect of tau on kinesin is well established *in vitro* for the ubiquitous kinesin heavy chain isoform 5B (Kif5B) (13, 15), we assayed the effect of a DOPC membrane on Kif5B-based transport in addition to the neuronal isoform Kif5A, mutations in which cause neurodegeneration (3-5).

**Table 1.**
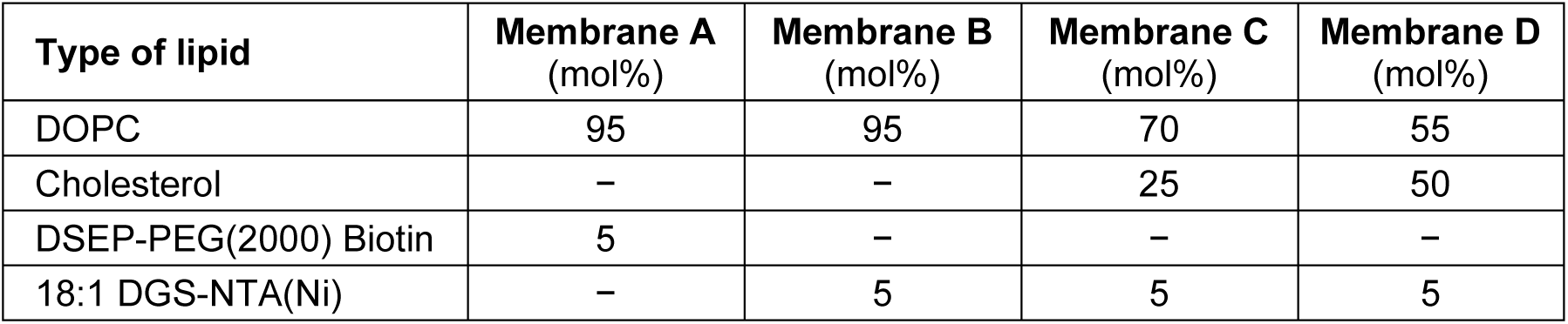
Membrane lipid compositions. DOPC: dioleoylphosphatidylcholine, a neutral lipid that models the fluid membrane of intracellular cargos (28, 29). DSEP-PEG(2000) Biotin: 1,2-distearoyl-sn-glycero-3-phosphoethanolamine-N-[biotinyl(polyethylene glycol)-2000] (ammonium salt), a biotinylated lipid that mediates the attachment of the kinesin-1 heavy chain isoform 5B (residues 1-560, C-terminal biotin tag). 18:1 DGS-NTA(Ni): 1,2-dioleoyl-sn-glycero-3-[(N-(5-amino-1-carboxypentyl)iminodiacetic acid)succinyl] (nickel salt), a nickel-loaded chelator lipid that mediates the attachment of the kinesin-1 heavy chain isoform 5A (full-length, C-terminal hexahistidine tag).

| Type of lipid | Membrane A<br>(mol%) | Membrane B<br>(mol%) | Membrane C<br>(mol%) | Membrane D<br>(mol%) |
| --- | --- | --- | --- | --- |
| DOPC | 95 | 95 | 70 | 55 |
| Cholesterol | – | – | 25 | 50 |
| DSEP-PEG(2000) Biotin | 5 | – | – | – |
| 18:1 DGS-NTA(Ni) | – | 5 | 5 | 5 |

**Figure 1.**
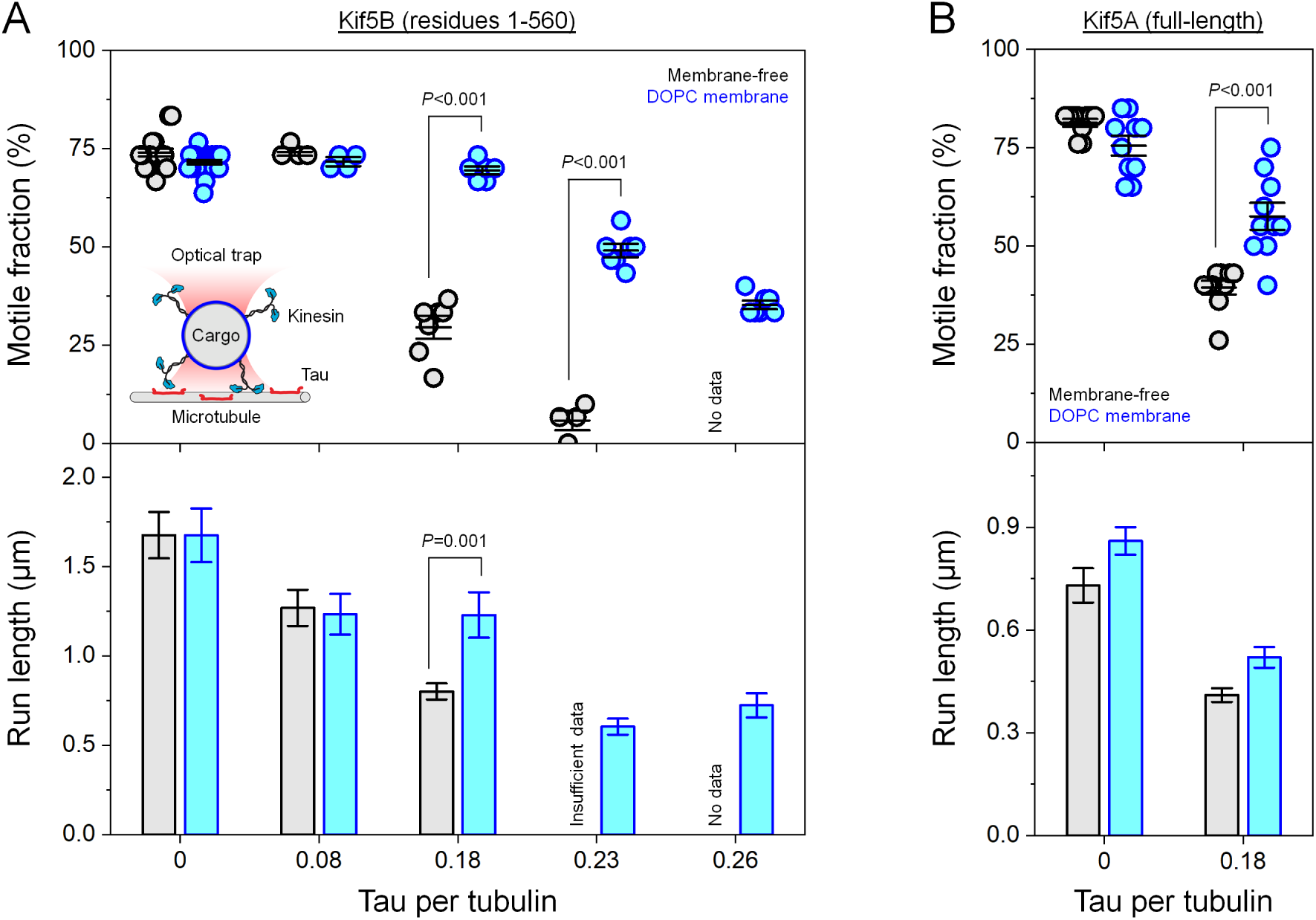
A fluid lipid membrane reduces the inhibitory effect of tau on kinesin-1. (*A*) Transport of membrane-free (grey) and membrane-enclosed (blue) cargos by the ubiquitous kinesin heavy chain isoform Kif5B, as a function of tau concentration (represented as the incubation ratio of tau per tubulin dimer). For each cargo type, the concentration of kinesin used to prepare the motor-cargo complex was empirically tuned to achieve the same motile fraction on tau-free microtubules (∼75%) and kept constant across all tau concentrations. *Inset*: Experimental schematic (not to scale). An optical trap confines an individual cargo near the microtubule. A motile event is scored if the cargo displays directed motion within 30 s. The trap is then shuttered to measure the zero-load run length. *Top*: Motile fraction of cargos. Horizontal lines indicate mean and 68% confidence interval. *n* = 4-22 trials, with 30 cargos measured in each trial. Significant *P*-values from two-sample Welch’s *t*-tests are indicated. *Bottom*: Mean run length of motile cargos. Error bars indicate standard error of the mean (SEM). *n* = 50-419 motile runs for the indicated mean run lengths. Low motile fractions of membrane-free cargos (<5%) at higher tau concentrations (0.23 and 0.26 tau per tubulin) resulted in insufficient data (*n* = 6 runs) or no data (*n* = 0 runs) for run-length determination. Mann-Whitney U tests returned one significant *P*-value. Run-length distributions are provided in *SI Appendix*, Fig. S2. (*B*) Cargo transport by the neuronal kinesin heavy chain isoform Kif5A, for two tau concentrations. Confidence intervals and error bars are as described in (*A*). *Top*: *n* = 10 trials, with 20 cargos measured in each trial. Two-sample Welch’s *t*-tests returned one significant *P*-value. *Bottom*: *n* = 66-191 motile runs. Run-length distributions are provided in *SI Appendix*, Fig. S3.

Attaching kinesins to a fluid lipid membrane reduced the inhibitory effect of tau on motile fraction for both the ubiquitous isoform Kif5B and the neuronal isoform Kif5A (Fig. 1, *Top*). We empirically tuned the concentration of motors in each cargo preparation to match the motile fraction for membrane-enclosed and membrane-free cargos in the absence of tau (∼75%; Fig. 1*A, Top*). This motile fraction corresponds to ∼30% of cargos being carried by two or more kinesins (34), mimicking the *in vivo* scenario (17-19). For the ubiquitous isoform Kif5B, the motile fraction of both membrane-free and membrane-enclosed cargos decreased with increasing concentrations of tau (Fig. 1*A, Top*). This result is consistent with prior reports that tau reduces the binding rate of kinesin for the microtubule (13-15). However, in contrast to membrane-free cargos (grey, Fig. 1*A, Top*), the motile fraction of membrane-enclosed cargos was less impacted by tau (blue, Fig. 1*A, Top*), suggesting that the binding rate of kinesin for tau-decorated microtubules was less impaired when the motors were membrane-bound. Similarly, for the neuronal isoform Kif5A, the motile fraction was significantly higher for membrane-enclosed cargos than membrane-free cargos in the presence of tau (blue, Fig. 1*B, Top*). To control for the possibility that the cargo membrane removes tau from the microtubule, we repeated motile-fraction measurements using microtubules pre-incubated with excess DOPC lipid vesicles. Consistent with reports that tau binds anionic but not neutral lipids (35), preincubating microtubules with neutral DOPC lipid vesicles did not impact motile-fraction measurements (*SI Appendix*, Fig. S1).

The effect of a fluid membrane on cargo run length differed between the two kinesin isoforms (Fig. 1, *Bottom*; and *SI Appendix*, Figs. S2 and S3). Here we confirmed that a fluid DOPC membrane did not impact cargo run length in the absence of tau (Fig. 1*A, Bottom*; and reference (23)). We also verified that the run lengths of both isoforms decreased in the presence of tau (Fig. 1, *Bottom*; and references (14, 15)). For the ubiquitous isoform Kif5B, cargo run-length was less impacted by tau when the motors were membrane-bound (Fig. 1*A, Bottom*). In contrast, for the neuronal isoform Kif5A, there was no significant difference in tau’s effects on the run length of membrane-enclosed cargos and membrane-free cargos (Fig. 1*B, Bottom*). A key difference between the isoforms is that Kif5A unbinds from the microtubule substantially faster than Kif5B (*SI Appendix*, Fig. S4). Although the run length of few-motor cargos is sensitive to changes in the binding rate of the motor (as reflected by the motile-fraction data in Fig. 1, *Top*), it is generally established that this sensitivity decreases as the motor’s unbinding rate increases (32, 33). Our run-length data (Fig. 1, *Bottom*) are consistent with this general relation between unbinding rate and the sensitivity of run length to binding rate. Hereafter, we focused our study on the neuronal isoform Kif5A based on its significance in neuronal physiology and human health (3-5).

### Membrane cholesterol amplifies the inhibitory effect of tau on kinesin-1

We next examined whether adding cholesterol to the DOPC bilayer impacts the transport of membrane-enclosed cargos. We added up to 50 mol% cholesterol to the DOPC lipid bilayer (Table 1), reducing membrane fluidity and motor diffusion by ∼3-fold (24, 27) without microdomain formation (36). The concentration of nickel-loaded chelator lipid was kept constant for identical Kif5A attachment across cholesterol levels (Table 1). In addition to the *in vivo* scenario of ∼1-2 kinesin per cargo examined in Figure 1, we assayed a higher motor number range that is anticipated (32) to increase the sensitivity of cargo run length to changes in the binding rate of the motor; a higher motor number range may also be relevant for long-range transport *in vivo* (24).

For cargos carried by ∼1-2 kinesins, adding cholesterol to the DOPC bilayer amplified the inhibitory effect of tau on motile fraction but not on run length (Fig. 2*A*). Here we employed the same Kif5A concentration as in Figure 1*B*. In the absence of tau, the fraction of motile cargos remained constant at ∼75% irrespective of membrane cholesterol; the mean run lengths of these motile cargos were also unaffected (Fig. 2*A*). These tau-free results are consistent with prior reports (23, 24). In the presence of tau, the fraction of motile cargos declined ∼2-fold further as membrane cholesterol increased to 50 mol% (Fig. 2*A, Top*), however, there was no further decrease in run length over this range of increasing cholesterol concentration (Fig. 2A, *Bottom*; and *SI Appendix*, Fig. S5). These results recapitulate our data comparing membrane-enclosed cargos with membrane-free cargos at the same Kif5A concentration (Fig. 1*B*), suggesting that changes in kinesin diffusion underlie both sets of observations. The significant effect of cholesterol on motile fraction (Fig. 2*A, Top*) indicates that cholesterol amplifies the inhibitory effect of tau on the binding rate of kinesin. The null effect of cholesterol on run length (Fig. 2*A, Bottom*) on the other hand indicates that the run length in these experiments is minimally impacted by the changes in binding rate. Furthermore, under conditions for which the cargo run length is insensitive to binding rate, current descriptions indicate that the run length of the cargo is primarily determined by the unbinding rate of the single motor (32, 33). Our run-length data here (Fig. 2*A, Bottom*) thus indicate that increases in membrane cholesterol do not significantly impact the unbinding rate of Kif5A.

**Figure 2.**
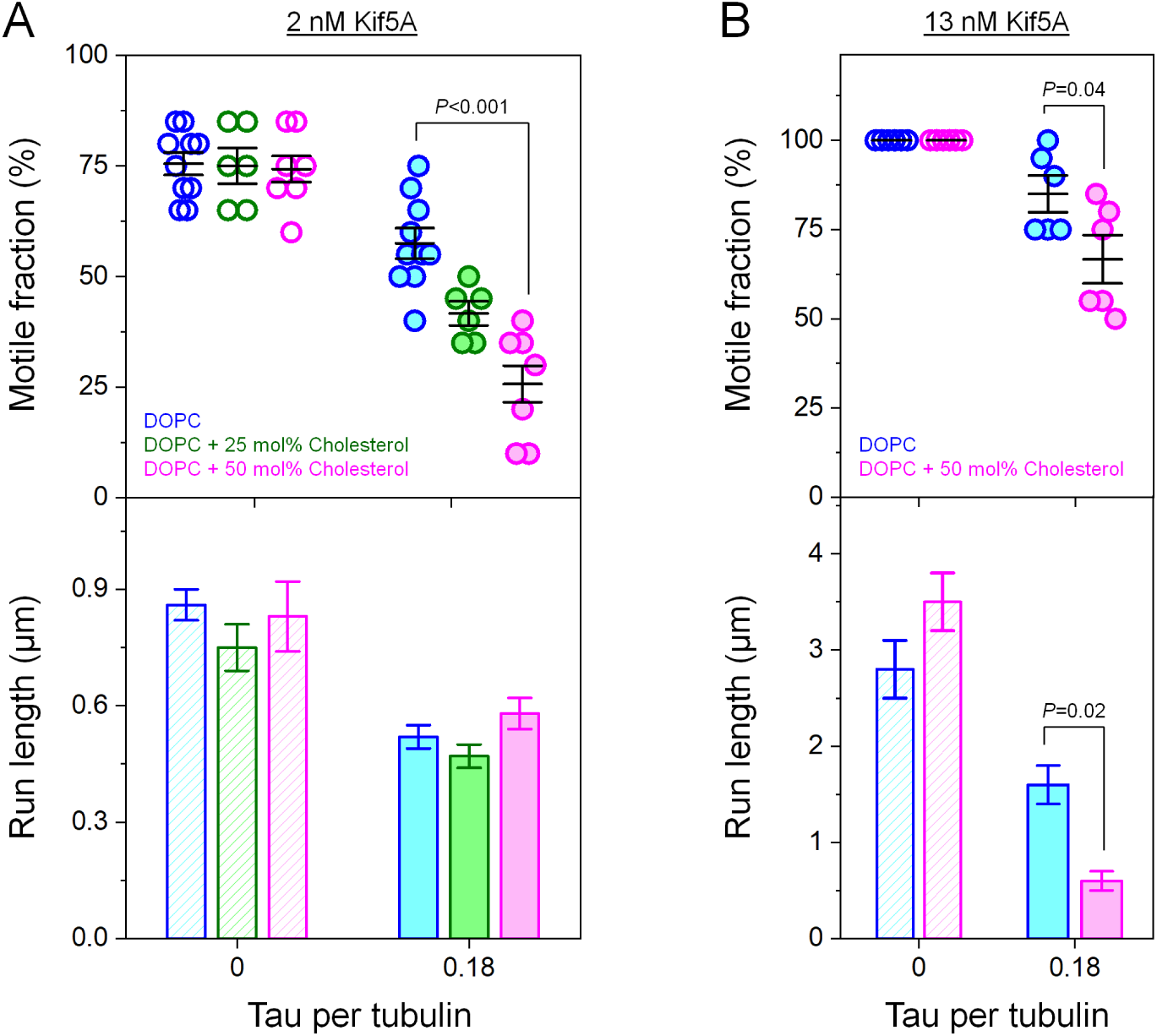
Membrane cholesterol amplifies the inhibitory effect of tau on kinesin-1. (*A*) Transport of membrane-enclosed cargos in the physiological range of ∼1-2 kinesins per cargo (as in Figure 1*B*) in the absence and presence of tau, for three cholesterol levels (blue: 0 mol%; green: 25 mol%; magenta: 50 mol%). The concentration of Kif5A was kept constant at 2 nM. *Top*: Motile fraction of cargos. Horizontal lines indicate mean and 68% confidence interval. *n* = 6-10 trials, with 20 cargos per trial. Kruskal-Wallis *H* tests returned one significant *P*-value. *Bottom*: Mean run length of motile cargos. Error bars indicate SEM. *n* = 21-113 motile runs. Run-length distributions are provided in *SI Appendix*, Fig. S5. (*B*) Transport of membrane-enclosed cargos at a higher motor number range (∼4 or more kinesins per cargo) in the absence and presence of tau, for two cholesterol levels (blue: 0 mol%; magenta: 50 mol%). The concentration of Kif5A was kept constant at 13 nM. Confidence intervals and error bars are as described in (*A*). *Top*: *n* =10 trials, with 20 cargos per trial. Two-sample Welch’s *t*-tests indicated one significant *P*-value. *Bottom*: *n* = 52-116 motile runs. Mann-Whitney U tests returned one significant *P*-value. Run-length distributions are provided in *SI Appendix*, Fig. S6.

For cargos carried by substantially more motors, membrane cholesterol amplified the inhibitory effect of tau on run length as well as on motile fraction (Fig. 2*B*). Here we increased the Kif5A concentration by ∼6-fold from that in Figure. 2*A*, resulting in run lengths >4-fold longer than that of cargos carried by a single Kif5A (*SI Appendix*, Fig. S4*A*). The extended run lengths indicate that the cargos were carried by ∼4 or more kinesins, based on prior results relating run length to kinesin motor number (33, 37-40). In the absence of tau, we again observed similar motile fractions and run lengths independent of membrane cholesterol (Fig. 2*B*). In the presence of tau, there was a further ∼2.6-fold decrease in cargo run length as membrane cholesterol increased from 0 to 50 mol% (Fig. 2*B, Bottom*; and *SI Appendix*, Fig. S6), in addition to a significant further decrease in motile fraction over the same cholesterol increase (Fig. 2*B, Top*). These results again indicate that cholesterol reduces the binding rate of kinesin in the presence of tau; they also highlight the potential of membrane cholesterol in impacting cargo run length even in the absence of a substantial effect on the unbinding rate.

### Membrane cholesterol reduces kinesin binding to microtubules in the presence of tau

We next expanded our experiments in Figure 2 to examine how membrane cholesterol impacts motile fraction over a broader range of the number of motors on the cargo (Fig. 3*A*). At each Kif5A concentration tested, membrane cholesterol consistently amplified tau inhibition of motile fraction (filled circles, Fig. 3*A*) but did not impact motile fraction in the absence of tau (open circles, Fig. 3*A*). At higher cholesterol levels, higher concentrations of Kif5A were needed to reach the same motile fraction on tau-decorated microtubules (Fig. 3*A*). The null effect of cholesterol in the absence of tau is in good agreement with prior reports that examined the binding of membrane-bound kinesins to tau-free microtubules (23, 24). The significant effect of cholesterol in the presence of tau demonstrates membrane cholesterol as a general and sensitive factor tuning the binding of kinesin-based cargos to tau-decorated microtubules.

**Figure 3.**
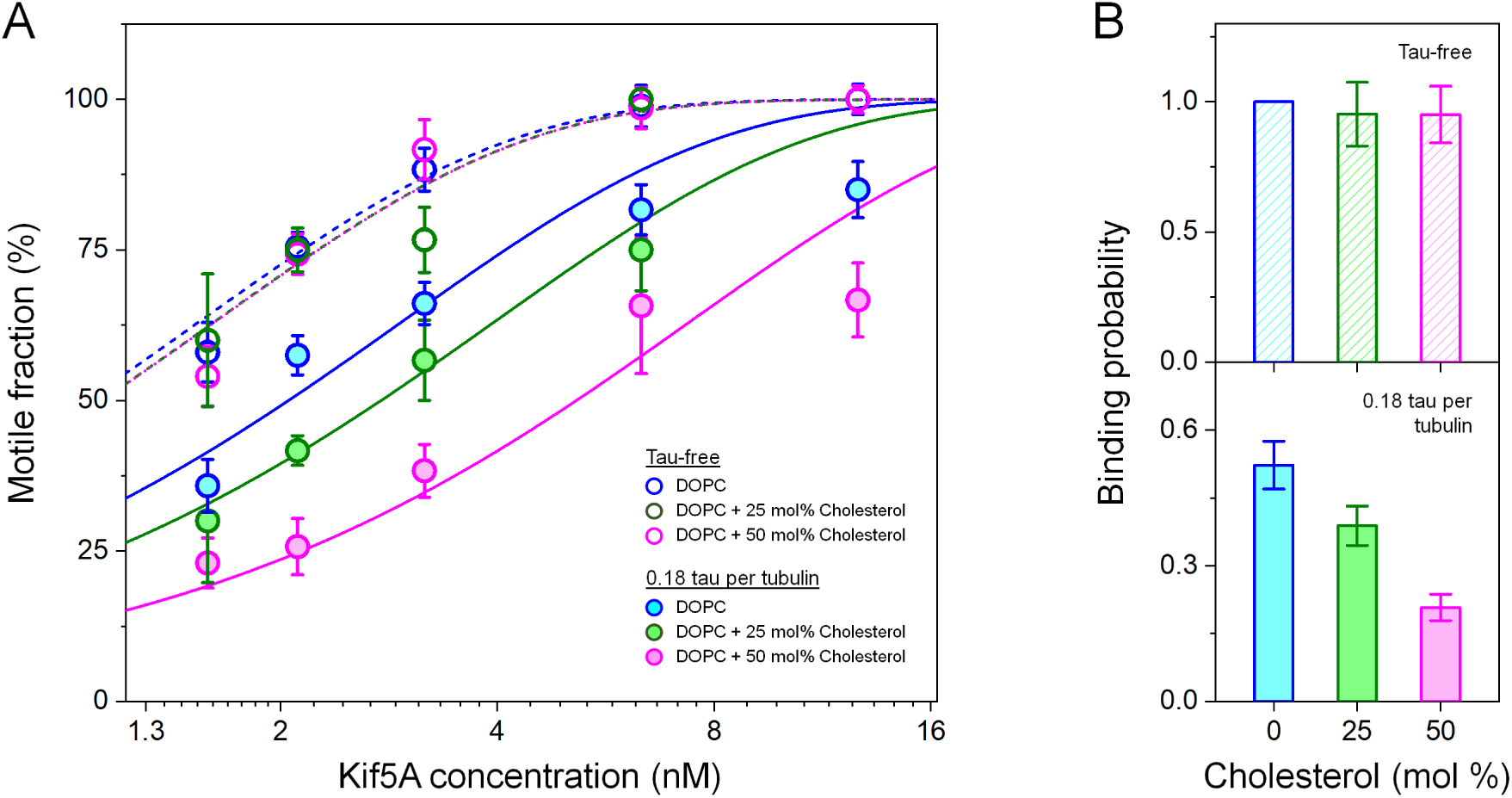
Membrane cholesterol reduces kinesin binding to microtubules in the presence of tau. (*A*) Motile fraction of membrane-enclosed cargos as a function of Kif5A concentration in the absence and presence of tau, for three cholesterol levels (blue: 0 mol%; green: 25 mol%; magenta: 50 mol%). Dashed and solid lines indicate best fits to our analytical model of motile fraction (Eqn. 1), with fitting parameter *α* shared among all six datasets; and binding probability *p* constrained as 1 for tau-free data with 0 mol% cholesterol and determined separately for the remaining five datasets. Dashed lines indicate best fits to data from tau-free conditions; solid lines indicate best fits to data measured in the presence of tau; 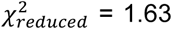. Error bars indicate SEM. *n* = 2-10 trials, with 20 cargos measured for each trial. Individual trials at 2 nM and 13 nM Kif5A are shown in Figure 2. (*B*) Microtubule-binding probability of Kif5A as a function of cholesterol levels, determined via the best-fits in (*A*). Error bars indicate the standard error of the fit.

We next derived an analytical model of motile fraction to determine how membrane cholesterol affects kinesin binding in the presence of tau. Motile fraction reflects both the number of motors available for transport and the probability that each motor binds the microtubule during the interaction time. Because we used the same concentration of motors to prepare the kinesin-cargo complexes for each set of comparisons, the number of motors on the cargo remained constant across cholesterol levels and tau concentrations. The observed reduction in motile fraction, both in the presence of tau and at higher cholesterol levels (Fig. 3*A*), therefore indicates reduction in the microtubule-binding probability of each motor on the cargo. Such reduction in binding probability, however, is not considered in the previous description of motile fraction (6, 30), which assumes that each motor binds to the microtubule with a probability of 1. Here we extended the analytical description of motile fraction to consider binding probabilities below 1. As in the previous derivation (6, 30), we considered the experimental scenario in which the number of motors on the cargo is Poisson-distributed. We then modeled motile fraction as the probability that each cargo is carried by at least 1 motor protein, multiplied by the probability that at least 1 of these motor proteins binds the microtubule, summed over all cargos in a population. Under the assumption that each motor binds the microtubule identically and independently, we have (*SI Appendix*, Note 1)

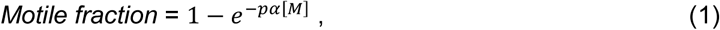

where 0 ≤ *p* ≤ 1 is the probability that a single motor binds the microtubule within the interaction time (30 s; Fig. 1*A, Inset*), and *α* is a fitting parameter relating the number of motors bound to the cargo to [*M*], the concentration of motors used to prepare the motor-cargo complex. As the binding probability decreases below 1, our model predicts that the fundamental dependence of motile fraction on motor concentration remains unchanged, whereas the motor concentration is effectively rescaled by the binding probability *p*.

Our model in Eqn. 1 accurately captures our motile-fraction measurements (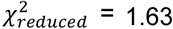; Fig. 3*A*). Because we employed the same motor-attachment strategy across experiments (Table 1), the fitting parameter *α* in Eqn. 1 was shared among datasets. Additionally, kinesin was also previously shown to have a unit probability of binding to tau-free microtubules within the interaction time of 30 s (14, 30, 31). We therefore constrained the binding probability for one set of the tau-free data (0 mol% cholesterol) to be 1 and examined how this binding probability is impacted by changes in tau concentration or cholesterol levels. In the absence of tau, adding cholesterol to the cargo membrane did not significantly impact the binding probability of kinesin, which remained 1 within fitting uncertainty (Fig. 3*B, Top*). In the presence of tau, the best fits returned binding probabilities below 1 (Fig. 3*B, Bottom*); there was a ∼2.5-fold further reduction in binding probability as membrane cholesterol increased from 0 mol% to 50 mol% (Fig. 3*B, Bottom*). Multiplying motor concentrations by these binding probabilities collapsed motile-fraction measurements onto a single master curve (*SI Appendix*, Fig. S7). These results indicate that the fundamental mechanism underlying motile probability is not impacted by tau or membrane cholesterol. Instead, reduction of motile fraction by tau and membrane cholesterol may be understood as effectively rescaling the kinesin concentration by the binding probability of the motor. Tau reduces the binding probability and membrane cholesterol amplifies tau inhibition, further reducing binding probability. Because binding probabilities were measured over the same interaction time (30 s; Fig. 1*A, Inset*), reduction in binding probability reflects reduction in the rate that the motor binds the microtubule. Our results thus indicate that membrane cholesterol reduces the rate that kinesin binds the microtubule in the presence of tau.

### A mechanistic model linking motor diffusion to kinesin binding in the presence of tau

We next developed a model describing how membrane cholesterol reduces kinesin binding in the presence of tau. Cholesterol reduces the diffusion of kinesin in the cargo membrane (24, 27) whereas tau occludes kinesin’s binding sites on the microtubule (10-12). We therefore considered a simple 1-dimensional model in which a motor diffusively searches for open binding sites on the microtubule (Fig. 4*A*). We assumed that open binding sites are separated by a characteristic distance Δ*x*; an increase in this characteristic distance recapitulates binding-site occlusion by tau. Note that, tau can diffuse on the microtubule and/or condense into islands that are kinetically distinct from monomeric tau (11, 12, 41, 42), these higher order effects on the spatial distributions were not included in the model. We further assumed that the motor binds each open binding site identically. Each time the motor encounters an open binding site, there is a likelihood *η* that the motor binds. If the motor does not bind at an encounter, then the motor diffuses to the nearest open site where it again has a likelihood *η* to bind. Under these simplifying assumptions, the rate that the motor encounters an open binding site is *D*/Δ*x*^2^ (*SI Appendix*, Note 2), the rate that the motor binds an open binding site is *Dη*/Δ*x*^2^, and the probability that the motor binds the microtubule within interaction time *T* is

**Figure 4.**
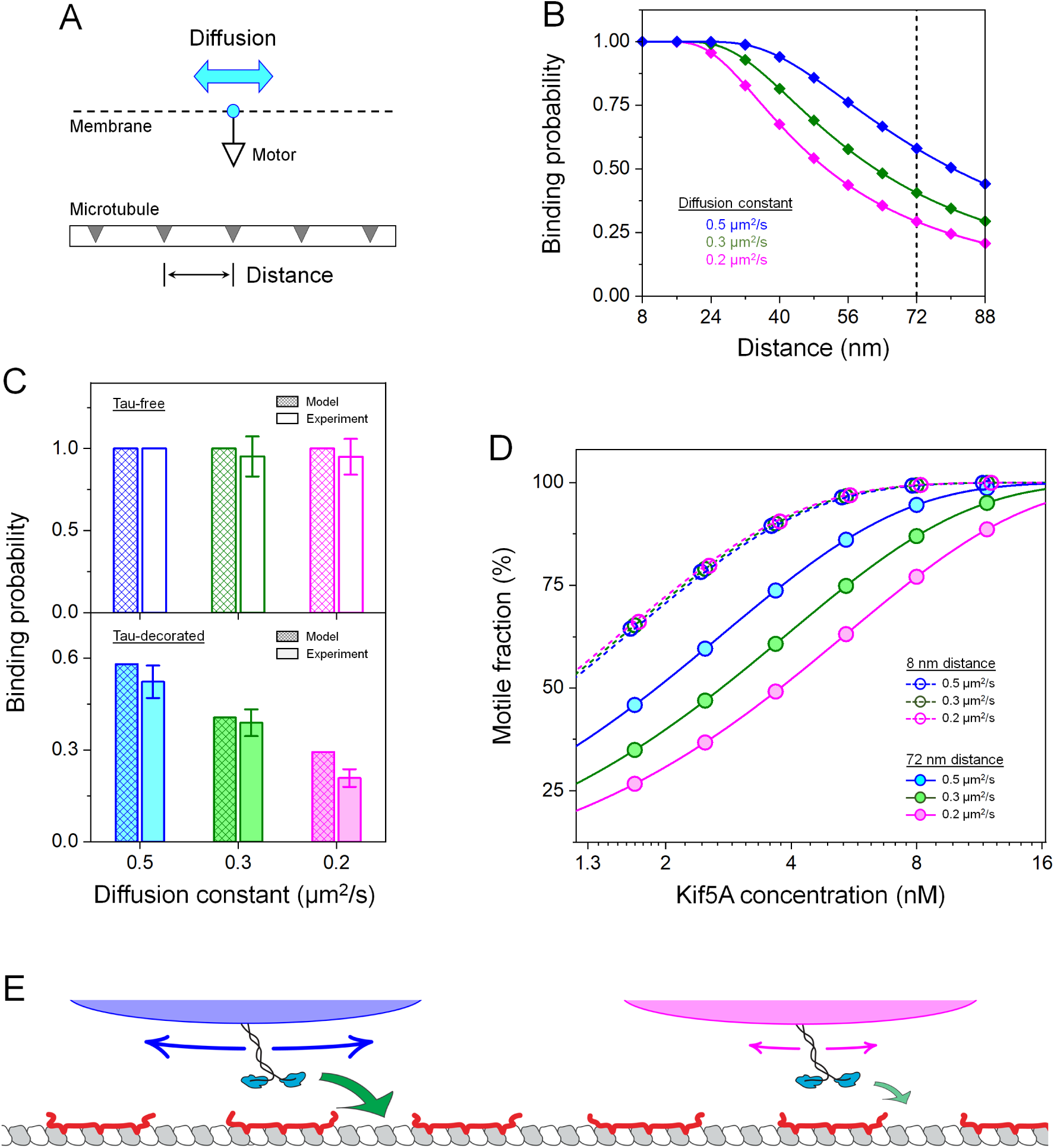
A mechanistic model linking motor diffusion to kinesin binding in the presence of tau. (*A*) Schematic of the model: A membrane-bound motor diffusively searches for open binding sites (shaded regions) on the microtubule. The motor and microtubule are vertically offset for visualization. The effect of tau is modeled as an increase in the characteristic distance between open binding sites on the microtubule. The effect of membrane cholesterol is modeled as a decrease in the diffusion constant of the motor in the cargo membrane. The analytical form of this one-dimensional model is shown in Eqn. 2. (*B*) Model predictions of the probability that a kinesin binds a microtubule within the interaction time of 30 s, for motor diffusion constants that approximate the effects of cholesterol in our experiments (blue: 0.5 µm^2^/s; green: 0.3 µm^2^/s; magenta: 0.2 µm^2^/s). For a binding-site distance of 72 nm (vertical dashed line), the predictions of Eqn. 2 yield a good match to experiments employing tau-decorated microtubules. (*C*) Comparison of model predictions and experimental values of the microtubule-binding probability for a single kinesin. Model predictions are as determined in (*B*) for characteristic distances of 8 nm (*Top*) and 72 nm (*Bottom*). Experimental values are as determined in Figure 3b for 0 mol % cholesterol (blue), 25 mol % cholesterol (green), and 50 mol % cholesterol (magenta). (*D*) Motile fraction calculated based on Eqn. 1, using binding probabilities predicted in (*C*) and the fitting parameter *α* determined in Figure 3*A*. Motile fractions for a characteristic distance of 8 nm (dashed lines and open symbols) are laterally offset for visualization. (*E*) Proposed mechanism: reduction in kinesin diffusion in the cargo membrane underlies the rate-limiting effect of cholesterol on kinesin binding in the presence of tau.

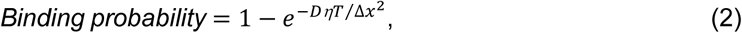

where *D* is the diffusion constant of the motor in the cargo membrane.

To compare our model with measurements, we estimated the variables in Eqn. 2 as follows. To estimate the diffusion constant *D*, we used the previously reported values of kinesin diffusion in DOPC membranes containing a similar range of membrane cholesterol (27) as a starting point, and reduced these values by 2.6-fold to account for the additional frictional drag (43) associated with the nickel-hexahistidine interaction in our Kif5A-membrane attachment strategy in accordance with previous results (44). We therefore estimated diffusion constants of 0.5, 0.3, and 0.2 µm^2^/s for Kif5A attached to DOPC membranes containing 0, 25, and 50 mol% cholesterol, respectively. For reference, the effective diffusion constant arising from the rotation of the membrane-free cargos (500-nm diameter) is ∼0.2 µm^2^/s. To estimate the binding likelihood *η* at each encounter, we combined our model for the binding rate of the motor (*Dη*/Δ*x*^2^) with a previous study that reported both the binding rate (4.7 ± 2.4 s^-1^) and the diffusion constant (1 ± 0.5 µm^2^/s) of a membrane-bound kinesin in the absence of tau (45). Assuming Δ*x* = 8 nm, the length of the individual tubulin dimer (46) and the step size of kinesin (47), as the distance between binding sites on tau-free microtubules, we estimated *η* = (3 ± 2)×10^−4^ as the likelihood that kinesin binds an open binding site at each encounter. Finally, we assumed that the motor is within reach of the microtubule for the full duration of *T* = 30 s that each cargo was held near a microtubule in our motile-fraction experiments (Fig. 1*A, Inset*). Thus, the only remaining variable in Eqn. 2 is the characteristic distance between open binding sites in the presence of tau.

Our simple model in Eqn. 2 captures the key observations in our experiments (Fig. 4 *B*-*D*). For each motor diffusion constant, Eqn. 2 predicts that the binding probability of the motor decreases as the characteristic distance between open binding sites increases (Fig. 4*B*). This prediction is in good agreement with our experiments employing increasing concentrations of tau (Fig. 1*A, Top*), indicating that increases in binding-site distance likely capture the key effect of tau in our experiments. As the motor diffusion constant decreased, the sensitivity of the binding probability to binding-site distance increased (Fig. 4*B*). At a characteristic distance of 8 nm, which reflects the tau-free condition, Eqn. 2 predicts a binding probability of 1 for all three diffusion constants (Fig. 4*B*), in good agreement with our experimental finding that membrane cholesterol does not impact kinesin-microtubule binding in the absence of tau (Fig. 4*C, Top*). At a larger characteristic distance of 72 ± 24 nm (uncertainty dominated by the estimate of *η* from previous measurements (45)), the predictions from Eqn. 2 match well with data from our experiments in the presence of tau (Fig. 4*C, Bottom*), capturing both the overall reduction in binding probability from the tau-free condition and the ∼2-fold further reduction in binding probability at higher cholesterol levels (Fig. 4*C, Bottom*). Based on recent structural evidence that each tau molecule binds to the microtubule longitudinally across three tubulin dimers (48), a characteristic distance of 72 ± 24 nm corresponds to an average of 3 ± 1 tau molecules between open binding sites. Note that, this characteristic distance is likely an overestimate, because the motor likely does not interact with the microtubule for the full duration that the cargo was held near a microtubule in our motile-fraction experiments. For example, the Brownian diffusion of the cargo in the optical trap may displace the motor beyond the interaction range with the microtubule. Combining these predicted binding probabilities with our description of motile fraction in Eqn. 1, we recapitulate the effects of cholesterol on motile fraction (Fig. 4*D*). Taken together, our simple model identifies diffusion of individual motors in the cargo membrane as a rate-limiting step in kinesin binding to microtubules in the presence of tau.

## Discussion

Intracellular cargos are often membrane-bound and transported by microtubule-based motors in the presence of MAPs. Here we established a direct link between the physical properties of the cargo membrane and MAP-based regulation of a major microtubule-based motor kinesin-1. Attaching kinesins to a fluid lipid membrane decreased the inhibitory effect of tau in comparison to membrane-free cargos (Fig. 1). Adding cholesterol to the cargo membrane increased the inhibitory effects of tau in a dosage-dependent manner (Figs. 2-3). These results reveal the potential of cargo membrane properties to regulate kinesin-based transport *in vivo*, for example under tau-enriched conditions such as in neurons, and/or in combination with factors (49-51) that alter tau binding to the microtubule.

We found that increasing the cholesterol level in the cargo membrane decreases the rate that kinesin binds microtubules in the presence of tau (Fig. 3*B*). Cholesterol is increasingly linked to cargo transport *in vivo* (19, 52, 53). The level of membrane-associated cholesterol in intracellular cargos can vary as part of the maturation process (19) or disease pathology (52, 53). Recent investigations have found that cholesterol-rich microdomains promote clustering of cytoplasmic dynein (19) and a monomeric kinesin (54) on the cargo membrane; such a clustering effect was not observed for the dimeric kinesin-1 (19). Here we employed levels of cholesterol within its solubility limit (∼67 mol% in DOPC, reference (36)), thus reducing membrane fluidity and motor diffusion (24, 27) without microdomain formation (36). Our findings therefore identify a new mechanism through which cholesterol can impact motor protein-based transport, which may be important for tuning motility without changing the number of kinesins carrying the cargo, for example in neuronal transport of cargos driven by a stable assembly of kinesins (17, 55).

We propose that reduction in kinesin diffusion in the cargo membrane underlies the rate-limiting effect of cholesterol on kinesin binding in the presence of tau (Fig. 4*E*). Our simple model (Eqn. 2) is akin to the collision theory of chemical reactions, which describes reaction rate as the product of the frequency of collision between reactants and the likelihood that each collision results in a reaction. Here we examined the binding rate between two reactants: the open binding sites on the microtubule and the individual kinesin molecule that diffuses in the cargo membrane. Tau reduces the frequency of collision between reactants by reducing the concentration of one reactant (the open binding sites), whereas motor diffusion tunes the time that it takes for one reactant (the unbound motor) to encounter the other reactant (the open binding sites) via diffusive search. An increase in motor diffusion constant decreases the diffusive search time for open binding sites and reduces inhibition by tau, whereas a decrease in motor diffusion constant increases the diffusive search time and amplifies inhibition by tau. *In vivo*, these membrane-based changes in binding rate may impact motile probability at lower tau concentrations, as the cargo likely has a shorter time to bind than in the experiments (30 s interaction time; Fig. 1*A, Inset*).

By necessity, our simple model in Eqn. 2 does not capture the full range of molecular details present in our experimental system. For example, we examined motor diffusion in one dimension, whereas the actual diffusion process occurs on the two-dimensional cargo surface. We also did not include details related to the distribution or the dynamics of tau on the microtubule (11, 12, 41, 42), the Brownian diffusion of the cargo, or how far the motor can extend to reach the microtubule at the cargo-microtubule interface (34, 56). Additionally, as the distance between binding sites and thus the diffusive search time decrease, molecular interactions between kinesin and individual binding sites (57-59) likely become rate-limiting for kinesin binding to the microtubule (as indicated by our tau-free data, Figs. 1-3; and references (23, 24)). Further work integrating these molecular details will help extend the model that we developed here to refine the mechanism underlying the kinesin-microtubule binding process.

The effects of membrane cholesterol may more broadly extend to other lipid alterations that impact motor diffusion in the cargo membrane, including those associated with aging and neurological diseases (20, 21). Similarly, factors other than MAP tau can occlude kinesin binding sites on the microtubule (13, 60-62). We anticipate that diffusion of individual motors in the cargo membrane may constitute a general mechanism influencing binding to the microtubule. Such a membrane-based mechanism may shed light on how age-related changes in lipid membranes might help explain why some inherited mutations in kinesins result in delayed onset diseases (such as those caused by Kif5A (3-5)). Conversely, our work suggests the possibility of targeting membrane fluidity to overcome transport dysfunctions that are a common early hallmark in aging and neurological diseases (22). Finally, extensive investigations have identified physiological factors that affect binding interactions between kinesin and tubulin, including MAPs (9, 63, 64) and post-translational modifications of tubulin (65-68). The work presented here provides a general framework for future studies examining the interplay between these factors and cargo membrane properties and how they contribute to the rich regulation of kinesin-based transport *in vivo*.

## Materials and Methods

### Materials

Human kinesin-1 heavy chain isoform Kif5A (full-length) with a C-terminal hexahistidine tag was expressed in *Escherichia coli* BL21(DE3) cells and purified as described (69); the final construct was a Kif5A homodimer with two C-terminal hexahistidine tags. Human kinesin-1 heavy chain isoform Kif5B consisting of residues 1-560 with a C-terminal Halo tag was expressed in *E. coli* BL21(DE3) cells and purified as described (23). The purified Kif5B protein was incubated with 10 µM HaloTag PEG-Biotin ligand (Promega) for 30 min on ice to produce a final construct of Kif5B homodimer with two C-terminal biotin tags. Unbound ligand was removed via dialysis and buffer was exchanged using 3000 Da dialysis membrane (Fisher). Human tau isoform encoding 352 amino acids with three microtubule-binding repeats (hTau23, or 3RS tau) was expressed in *E. coli* and purified as described (14). Tubulin was purified from bovine brain via two cycles of polymerization and depolymerization and phosphocellulose chromatography as described (14). All proteins (Kif5A, Kif5B, tau, and tubulin) were flash frozen and stored at −80 °C until use. Lipids 18:1 (Δ9-Cis) PC (DOPC), 18:1 DGS-NTA(Ni), and DSEP-PEG(2000) Biotin were purchased from Avanti Polar Lipids, Inc. Cholesterol (C_27_H_46_O) was purchased from Sigma-Aldrich. Streptavidin was purchased from Thermo Fisher. Chemicals were purchased from Sigma-Aldrich. Silica microspheres (0.5 µm in diameter; 10.2% solids) were purchased from Bangs Laboratories. Carboxylated polystyrene microspheres (0.5 µm in diameter; 2.5% solids) were purchased from Polysciences.

### Membrane-enclosed cargo preparation

Membrane-enclosed cargos were prepared as described (23, 25, 26) with modifications. Briefly, lipid mixtures (Table 1) were dried under vacuum at room temperature overnight, resuspended to 4 mM in HNa100 buffer (pH 7.0, 20 mM HEPES, 100 mM NaCl, 1 mM EGTA, 1 mM dithiothreitol) via five freeze-thaw cycles, and passed through a 30-nm polycarbonate membrane 21 times using a mini-extruder (Avanti Polar Lipids) to generate small unilamellar vesicles. Silica microspheres (40 µL stock) were bath-sonicated in 1 mL methanol for 1 min, resuspended in 1 mL 1 N KOH and bath-sonicated in ice water for 10 min, washed seven times in 1 mL Nanopure water, and resuspended in 40 µL Nanopure water. Freshly washed silica beads (9 µL) were incubated with 150 µL 2 mM freshly extruded small unilamellar vesicles for 30 min at room temperature, washed four times in 1 mL HNa100, resuspended in 100 µL 5 mg/mL casein solution in PMEE buffer (35 mM PIPES, 5 mM MgSO_4_, 1 mM EGTA, 0.5 mM EDTA, pH 7.1) for blocking at room temperature for 30 min. For preparations using lipid mixtures containing a nickel-loaded chelator lipid (Membranes *B*-*D*, Table 1), the resulting membrane-enclosed cargos were kept at 4 °C for use within two days. For preparations using a lipid mixture containing a biotinylated lipid (Membrane *A*, Table 1), the membrane-enclosed cargos were washed three times in 1 mL HNa100, resuspended in 40 µL 1 mg/mL streptavidin at room temperature for 1 h, washed three times in 1 mL HNa100, resuspended in 100 µL HNa100, and kept at 4 °C for use within two days.

### Lipid vesicle preparation

A cholesterol-free DOPC lipid mixture (Membrane *B*, Table 1) was dried under vacuum and resuspended in HNa100 as described above for membrane-enclosed cargos. The lipid mixture was then extruded through a 100-nm polycarbonate membrane 21 times using a mini-extruder (Avanti Polar Lipids). Lipid vesicles were used immediately following extrusion.

### Kinesin/cargo complex preparation

Kif5A was attached to membrane-enclosed cargos via a nickel-hexahistidine interaction (23, 28). Kif5B was attached to membrane-enclosed cargos via a streptavidin-biotin interaction (24, 26, 28). Both kinesin isoforms were attached to membrane-free polystyrene microspheres via non-specific interactions (23, 31, 34). Kinesin motors (1.5-12.6 nM Kif5A or 1.6-4.5 nM Kif5B) were incubated with membrane-enclosed cargos (3.3×10^5^ particles/µL) or membrane-free cargos (3.6×10^5^ particles/µL) in motility buffer (67 mM PIPES, 50 mM CH_3_CO_2_K, 3 mM MgSO_4_, 1 mM dithiothreitol, 0.84 mM EGTA, 10 µM taxol, pH 6.9) for 10 min at room temperature. The solution was then supplemented with an oxygen-scavenging solution (250 µg/mL glucose oxidase, 30 µg/mL catalase, 4.6 mg/mL glucose) and 1 mM ATP, followed by immediate use for optical trapping experiments.

### Microtubule preparation

Tubulin (40 *μ*M) was incubated in PM buffer (100 mM PIPES, 1 mM MgSO_4_, 2 mM EGTA, pH 6.9) supplemented with 1 mM GTP for 20 min at 37 °C. The assembled microtubules were incubated with an equal volume of PM buffer supplemented with 40 *μ*M taxol and 1 mM GTP for 20 min at 37°C. Taxol-stabilized microtubules were diluted to 500 nM in microtubule buffer (PMEE buffer supplemented with 10 µM taxol and 1 mM GTP) and incubated with 0-130 nM hTau23 in microtubule buffer for 20 min at 37 °C, followed by immediate use for flow cell preparation. The concentration of tau in each microtubule preparation is represented as the incubation ratio of tau molecules per tubulin dimer. The inhibitory effect of tau on kinesin was verified by the decrease in motile fraction on tau-decorated microtubules. For each preparation, the inhibitory effect of tau on motile fraction remained constant (within measurement uncertainty) for the duration of each set of experiments (up to ∼5 h).

### Flow cell preparation

Flow cells were constructed by sandwiching a cover glass and a microscope slide using double-sided tape. The coverslip surface was plasma cleaned and coated with poly-L-lysine for microtubule immobilization (60). Freshly prepared microtubules (tau-free or tau-decorated) were diluted to 45 nM in microtubule buffer and introduced into flow cells for incubation at room temperature for 10 min. The flow cell was then rinsed with microtubule buffer and then blocked with 5.55 mg/mL casein in microtubule buffer for 10 min at room temperature. In experiments in which the microtubules were incubated with excess DOPC lipid vesicles, flow cells were further incubated with 0 or 100 µM freshly extruded lipid vesicles in 5.55 mg/mL casein solution for 5 min at room temperature. The flow cells were then rinsed and blocked with 5.55 mg/mL casein in microtubule buffer for 10 min at room temperature. All flow cells were used for optical trapping experiments within the same day of preparation.

### *In vitro* optical trapping experiments

*In vitro* optical trapping experiments were carried out in flow cells at room temperature as described (23, 31, 34). Briefly, freshly prepared kinesin-cargo complexes were introduced to flow cells with immobilized microtubules. A cargo diameter of 500 nm was used, this cargo size is well-suited for optical trapping and is in the range of vesicle sizes *in vivo* (17-19). A single-beam optical trap similar to that described previously (30) was used to position individual cargos near a microtubule. The trap power was limited to <20 mW at fiber output, such that the trap positioned individual cargos but did not stall cargos carried by a single kinesin. A motile event was scored if the cargo moved directionally away from the trap center within 30 s (14, 30, 31). The optical trap was then shut off to either release the unbound cargo or to measure cargo run length under zero external load. Cargo trajectories were imaged at 100× magnification via differential interference contrast microscopy and video-recorded using a Giga-E camera at 30 Hz. Each preparation of kinesin-cargo complexes was used to measure transport at two tau concentrations. Each preparation of microtubules was used to measure the transport of two or more cargo types. The measurement order of each experimental condition was shuffled to eliminate potential artifacts.

## Data analysis

Cargos were particle-tracked to 10 nm resolution (1/3 pixel) via a template-matching algorithm (70). The run length of each motile cargo was determined as the net displacement of the cargo along the microtubule between binding to and unbinding. The distribution of run lengths for each experimental condition was fitted to a single exponential decay. To account for the time that elapsed before the manual shutoff of the optical trap, only runs >250 nm were analyzed. Runs that exceeded the experimental field of view were also excluded from the fit. Mean run length and SEM for each distribution were determined as the fitted decay constant and uncertainty, respectively. Fitting the cumulative probability distribution of the run lengths, which does not require data binning, returned similar values of mean run lengths. The unbinding rate of the single kinesin (*SI Appendix*, Fig. S4) was determined as the ratio of the mean run length to the mean velocity of single-kinesin cargos; the associated SEM was determined via standard error propagation.

## Model derivation

Derivations of Eqns. 1-2 are detailed in *SI Appendix*, Notes 1-2.

## Statistical analysis

Two-sample Welch’s *t*-test was used to determine the statistical significance of differences between two distributions of motile-fraction measurements. The Mann-Whitney U test was used to determine the statistical significance of differences between two distributions of run-length measurements. The Kruskal-Wallis *H* test, a non-parametric analysis of variance (ANOVA), was used to determine the statistical significance of differences between multiple distributions of motile-fraction measurements or run-length measurements.

## Acknowledgments

This work was supported by the National Institutes of Health (Grant Nos. R15 GM120682 to J.X. and R01 NS048501 to S.J.K.) and the National Science Foundation (Grant No. ENG-1563280 to M.V.). This work was supported in part by the intramural research program of the National Heart, Lung, and Blood Institute, National Institutes of Health, Department of Human Services (K.C.N). We thank John A. Hammer, Serapion Pyrpassopoulos, Matthew J. Bovyn, Steven P. Gross, and Yasuharu Takagi for discussion. We thank Tiffany J. Vora for manuscript editing.

## Author Contributions

J.X. designed research; Q.L., J.X., and L.A.C. performed research; Q.L., J.X., and J.O.W. analyzed data; J.X., K.C.N, J.T.F., and J.S. developed models; W.Q., M.V., and S.J.K. provided proteins; J.X. wrote the paper; all authors edited the paper.

## Supplementary Information for

### Supplementary Text

**Note 1. Derivation of motile fraction model (Eqn. 1 in the main text)**. We considered the experimental scenario that the number of motors on a cargo follows a Poisson distribution (1, 2). We modeled motile fraction as the weighted sum of the probability that a cargo is carried by at least 1 motor protein, and the probability that at least 1 of these motor proteins binds the microtubule. We assumed that each motor on the cargo binds the microtubule identically and independently. For each preparation of motor-cargo complexes, the average number of motors on the cargo is *α*[*M*], where *α* is a fitting parameter determined by the motor-cargo attachment strategy, and [*M*] is the concentration of motors that we used to prepare the motor-cargo complex. The probability that the cargo carries *k* = 1, 2, 3, … motor(s) is determined by the Poisson probability

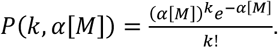

The probability that at least 1 of these *k* motors on the cargo binds the microtubule is 1 − (1 − *p*)^*k*^, where 0 ≤ *p* ≤ 1 is the probability that each motor binds the microtubule within the motile-fraction measurement time. Summing over all *k* values, we have

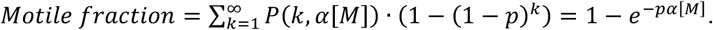

For *p* = 1, our model returns the same expression as that previously developed (1, 2), 1 − *e*^−*α*[*M*]^.

**Note 2. Derivation of the rate that a motor encounters an open binding site (Fig. 4 and Eqn. 2 in the main text)**. We modeled the microtubule lattice as a one-dimensional track with periodic binding sites. We then coarse-grained diffusion along this track to derive the time that a motor spends at each binding site before reaching an adjacent binding site. The inverse of this time is the rate that the motor encounters a new binding site. Specifically, we defined each binding site as an open interval of a width much smaller than the periodicity. Under this coarse-graining, the motor begins at the center of the binding site; the encounter ends when the motor diffuses to the next binding site centered ±Δ*x* away, where the process repeats itself. Solving the one-dimensional diffusion equation with the initial position of the motor at *x* = 0 and two absorbing boundaries at ±Δ*x*, we have

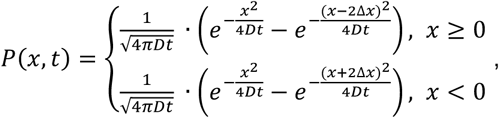

where *D* is the motor’s diffusion constant, and *P*(*x, t*) is the probability density of finding the motor at position *x* at time *t*. The probability that the motor remains within both boundaries for all times up to time *t* is

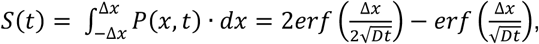

the probability that the motor first reaches either absorbing boundaries at exactly time *t* is

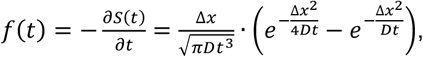

and the mean first-passage time of the motor to reach an adjacent binding region is

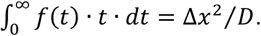

The inverse of this mean first passage time is the rate that the motor encounters an open binding site, *D*/Δ*x*^2^.

## Supplementary Figures

**Fig. S1.**
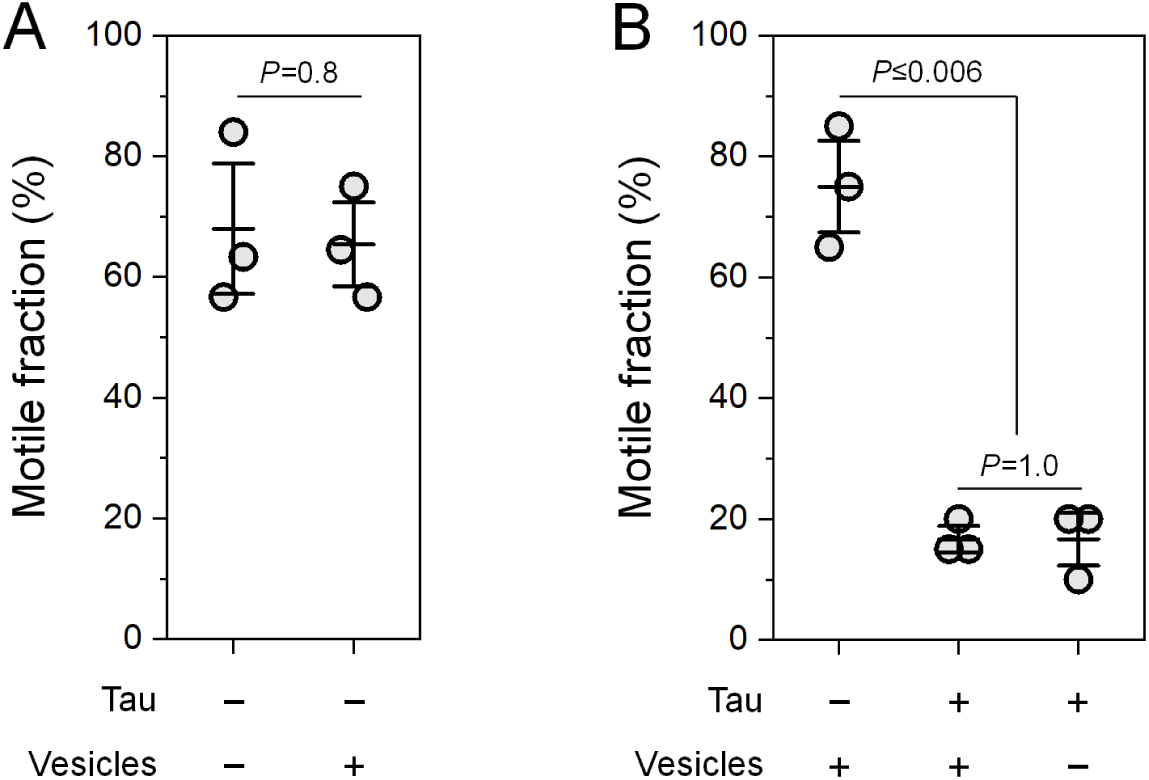
Preincubating microtubules with motor-free DOPC lipid vesicles does not impact cargo motile fraction. To control for the possibility that the cargo membrane binds and removes tau from the microtubule, microtubules were incubated with excess DOPC lipid vesicles prior to motile-fraction measurements (Materials and Methods in the main text). Membrane-free cargos were tested in order to probe potential membrane effects from vesicle preincubation alone. The concentration of Kif5A was kept constant in order to enable comparisons across experimental conditions. (*A*) Preincubating tau-free microtubules with 100 µM DOPC lipid vesicles (+Vesicles) did not impact cargo motile fraction. Horizontal lines indicate mean and 68% confidence interval. *n* = 3 trials, with 30 cargos measured in each trial. *P*-values from two-sample Welch’s *t*-tests are indicated. (*B*) Tau inhibition of motile fraction was not affected by preincubating microtubules with 100 µM DOPC lipid vesicles (+Vesicles). Tau-decorated microtubules (+Tau) were prepared at an incubation ratio of 0.18 tau per tubulin dimer. Confidence intervals, sample size, and statistical tests are as described in (*A*). These results are consistent with reports that tau binds anionic lipids but not neutral lipids (3) (such as the neutral DOPC lipid used here, Materials and Methods in the main text).

**Fig. S2.**
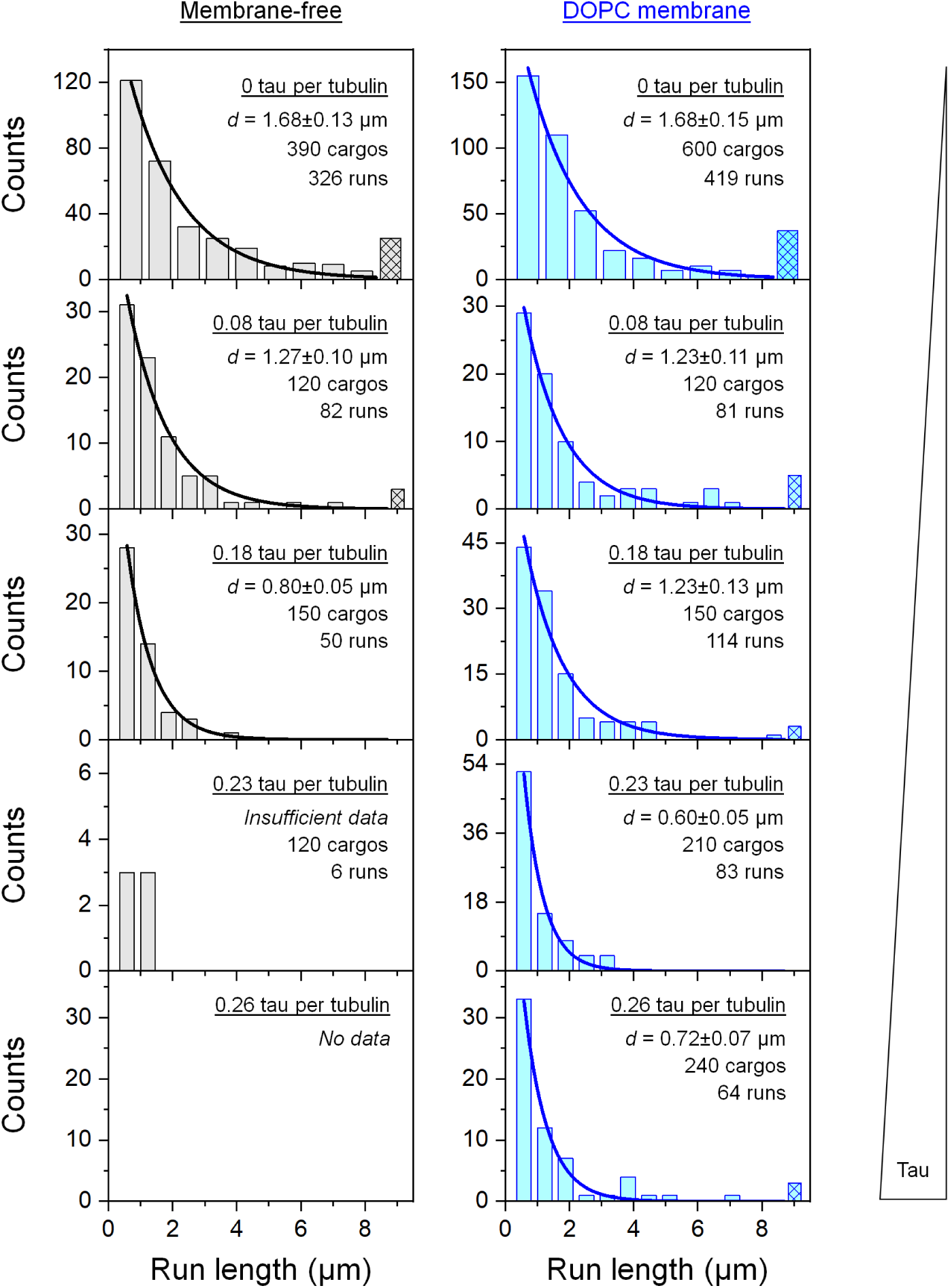
Run-length distributions of cargos carried by the kinesin heavy chain isoform 5B (Kif5B), corresponding to Figure 1*A* in the main text. Tau concentration, mean run length (*d*, ± standard error of mean (SEM)), number of cargos tested, and number of motile runs are indicated. Solid lines indicate single exponential fits to runs within the experimental field of view. Hatched bars indicate runs that exceeded the field of view.

**Fig. S3.**
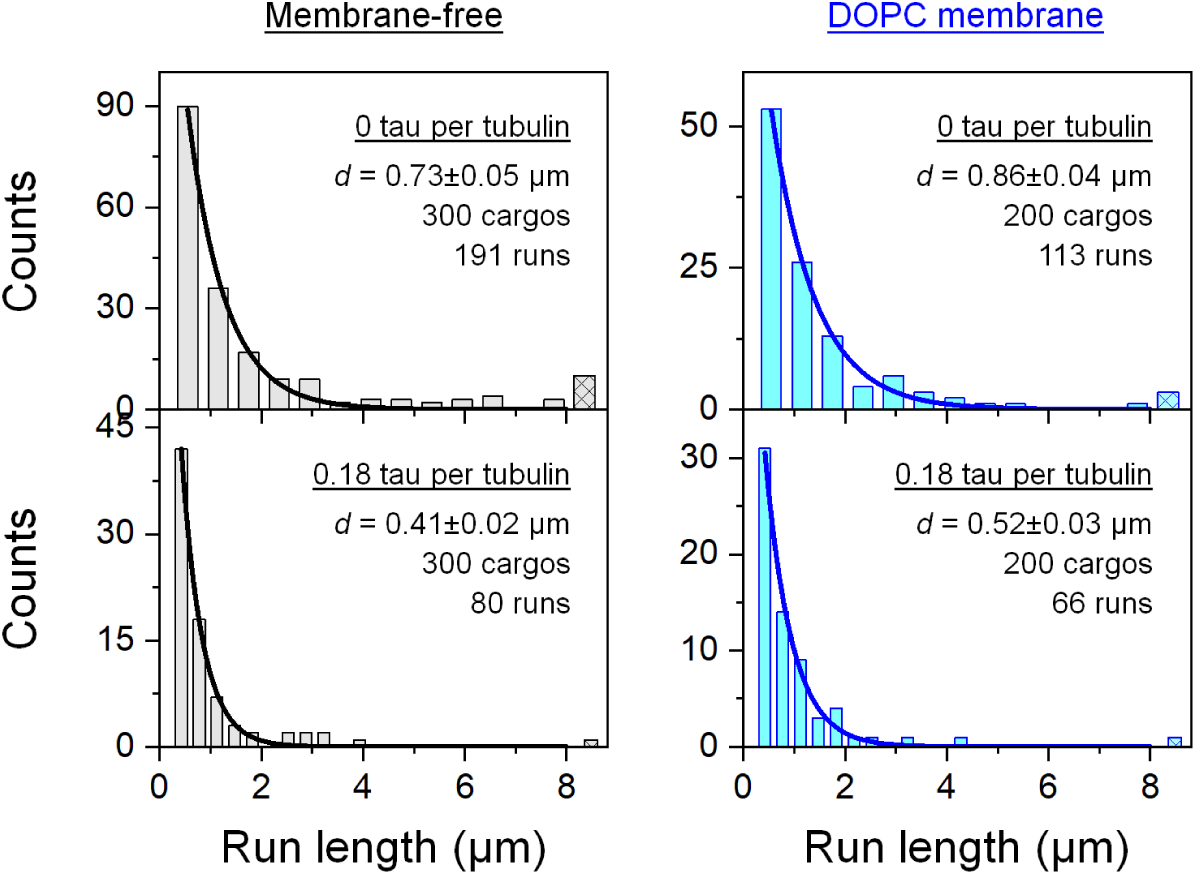
Run-length distributions of cargos carried by the kinesin heavy chain isoform 5A (Kif5A), corresponding to Figure 1*B* in the main text. Tau concentration, mean run length (*d*, ± SEM), number of cargos tested, and number of motile runs are indicated. Solid lines indicate single exponential fits to runs within the experimental field of view. Hatched bars indicate runs that exceeded the field of view.

**Fig. S4.**
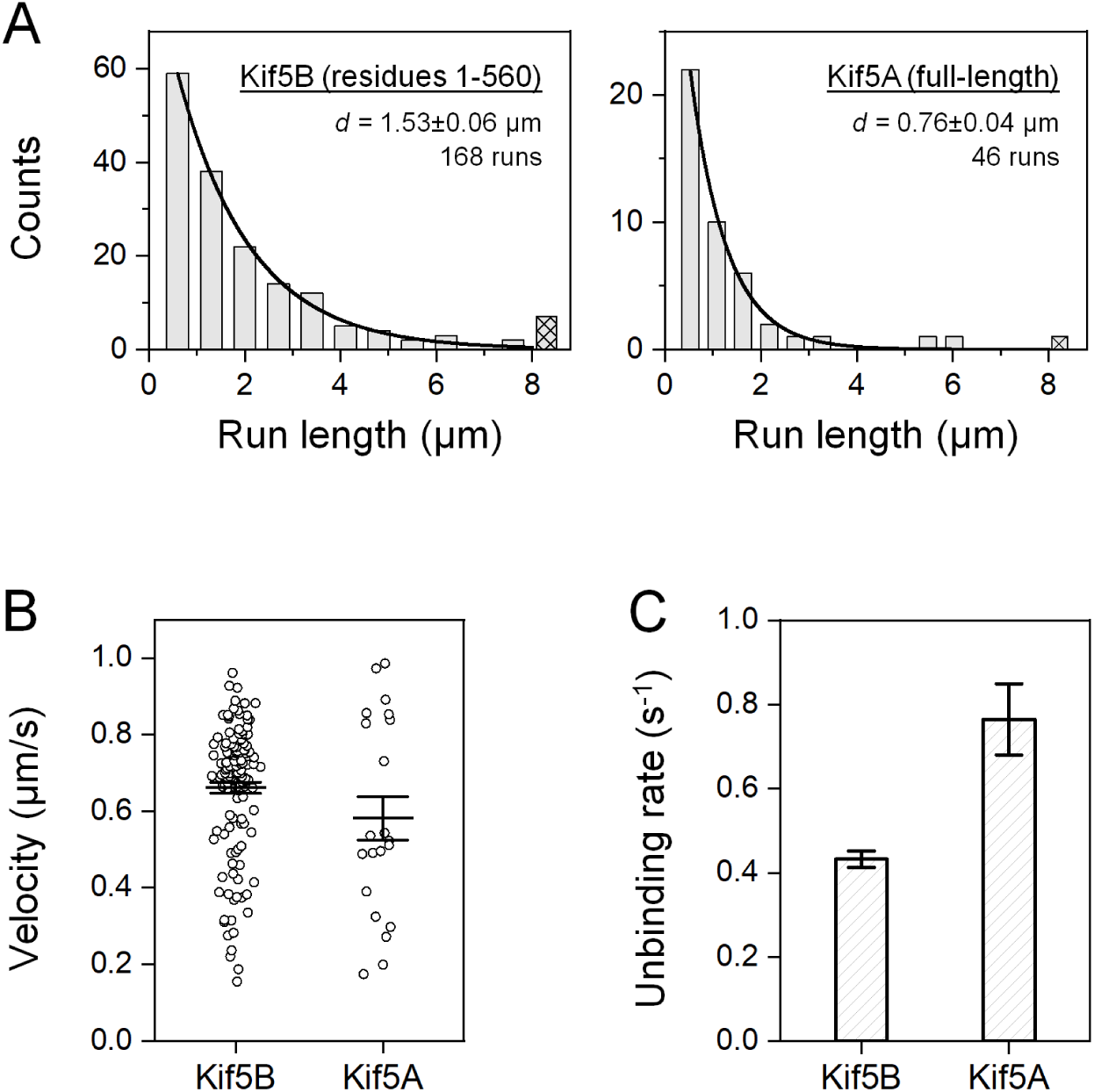
The neuronal isoform Kif5A unbinds from the microtubule substantially faster than the ubiquitous isoform Kif5B. (*A*) Run-length distributions of membrane-free cargos carried by a single Kif5B or Kif5A in the absence of tau. The concentration of kinesin was tuned such that the fraction of motile cargos in the absence of tau was ∼30%, indicating transport by a single motor (1, 2). Mean run length (*d*, ± SEM), number of cargos tested, and number of motile runs are indicated. Solid lines indicate single exponential fits to runs within the experimental field of view. Hatched bars indicate runs that exceeded the field of view. (*B*) Transport velocity of a single kinesin in the absence of tau. Horizontal lines indicate mean and 68% confidence interval. *n* = 139 and 21 runs. (*C*) Unbinding rate of a single kinesin from the microtubule in the absence of tau, determined as the ratio of the mean velocity (*B*) to the mean run length (*A*) for each isoform. Error bars indicate SEM. Kif5A unbinds ∼1.8 ± 0.2-fold faster than Kif5B.

**Fig. S5.**
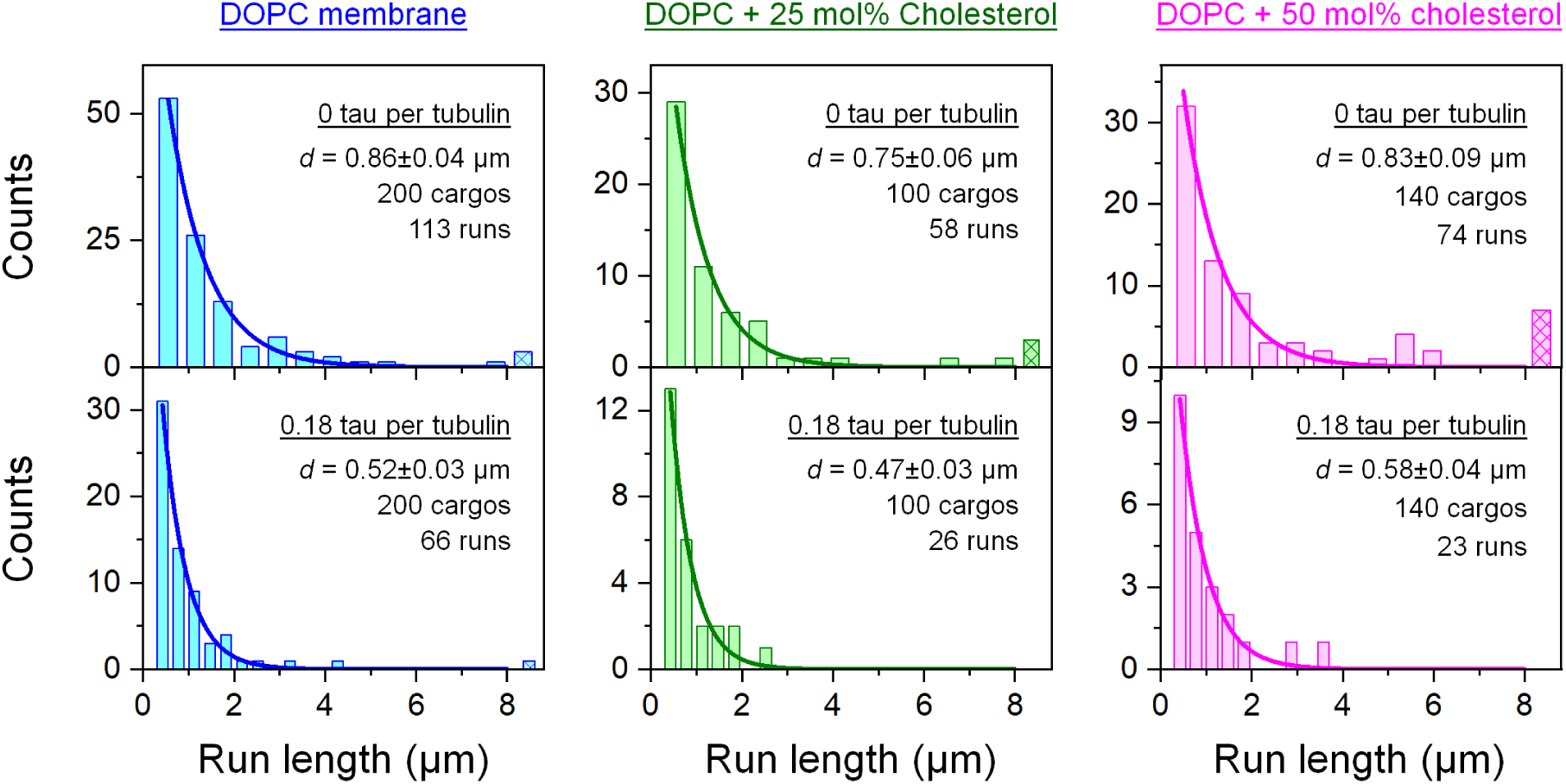
Run-length distributions of cargos carried by the neuronal isoform Kif5A, corresponding to Figure 2*A* in the main text. Tau concentration, mean run length (*d*, ± SEM), number of cargos tested, and number of motile runs are indicated. Solid lines indicate single exponential fits to runs within the experimental field of view. Hatched bars indicate runs that exceeded the field of view.

**Fig. S6.**
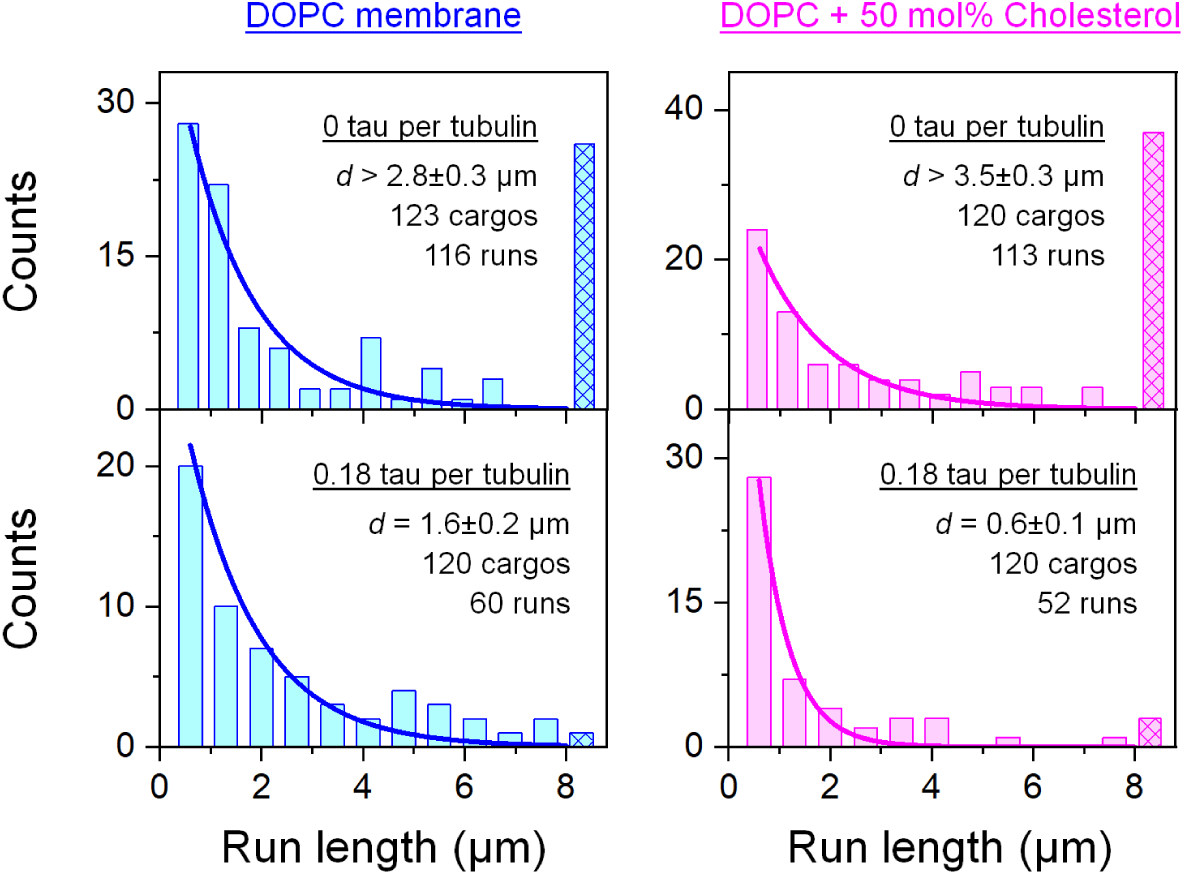
Run-length distributions of cargos carried by the neuronal isoform Kif5A, corresponding to Figure 2*B* in the main text. Tau concentration, mean run length (*d*, ± SEM), number of cargos tested, and number of motile runs are indicated. Solid lines indicate single exponential fits to runs within the experimental field of view. Hatched bars indicate runs that exceeded the field of view.

**Fig. S7.**
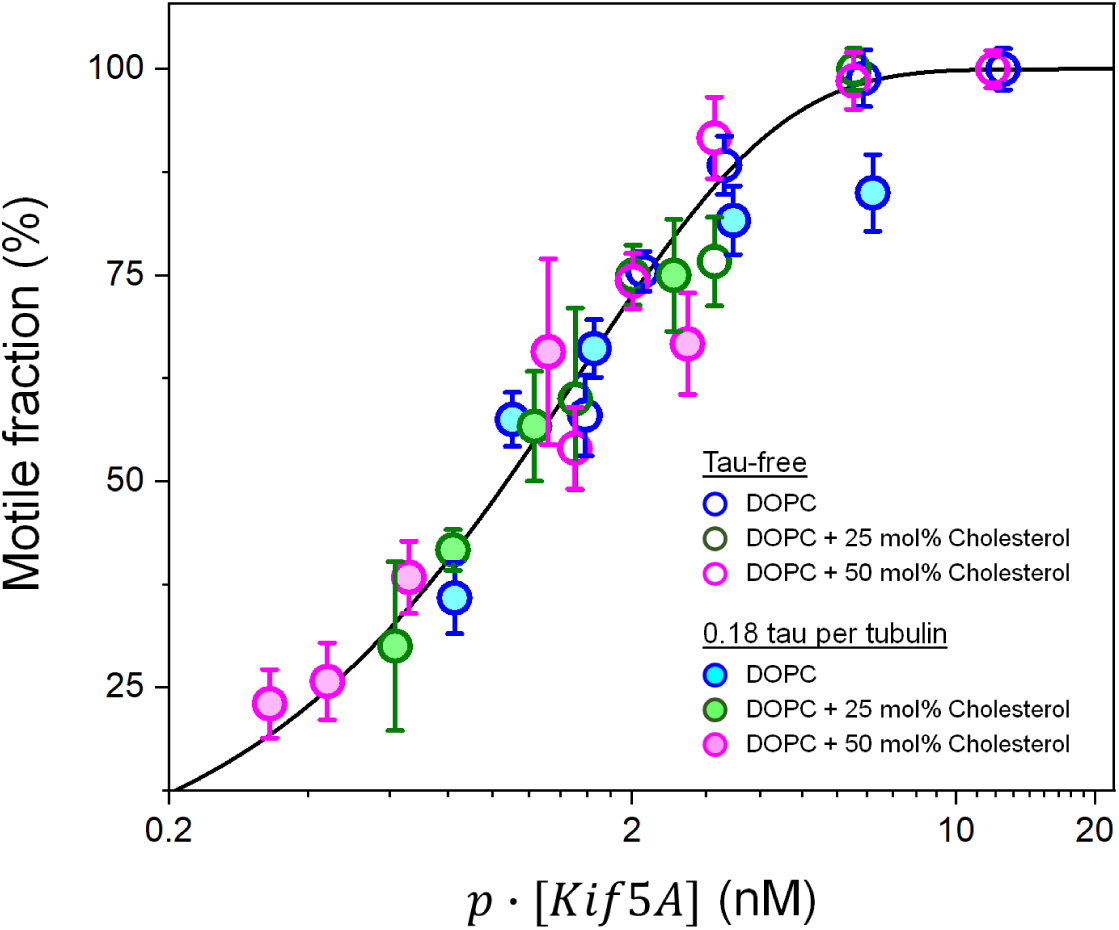
Multiplying kinesin motor concentration by binding probability collapses motile fraction measurements (open and filled circles) onto a single master curve (solid line). Motile fraction measurements, kinesin motor concentrations· [*Kif5A*], and binding probabilities *p* are as shown in Fig. 3 in the main text. Error bars indicate SEM. *n* = 2 - 10 trials, with 20 cargos measured for each trial. Solid line indicates the global fit to 1 − *e*^−*α*.*p*.[*KiK5A*]^, where *α* is a fitting parameter shared among datasets; 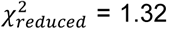. These results indicate that the fundamental mechanism underlying motile probability is not impacted by tau or membrane cholesterol. Instead, reduction of motile fraction by tau and membrane cholesterol may be understood as effectively rescaling the kinesin concentration by the binding probability of the motor. Tau reduces the binding probability and membrane cholesterol amplifies tau inhibition, further reducing binding probability (Fig. 3*B* in the main text).

## References

1. R. D. Vale, T. S. Reese, M. P. Sheetz, Identification of a novel force-generating protein, kinesin, involved in microtubule-based motility. Cell 42, 39–50 (1985).

2. S. T. Brady, A novel brain ATPase with properties expected for the fast axonal transport motor. Nature 317, 73–75 (1985).

3. E. Reid et al., A kinesin heavy chain (KIF5A) mutation in hereditary spastic paraplegia (SPG10). Am. J. Hum. Genet. 71, 1189–1194 (2002).

4. Y. T. Liu et al., Extended phenotypic spectrum of KIF5A mutations: From spastic paraplegia to axonal neuropathy. Neurology 83, 612–619 (2014).

5. A. Nicolas et al., Genome-wide Analyses Identify KIF5A as a Novel ALS Gene. Neuron 97, 1268–1283.e1266 (2018).

6. S. M. Block, L. S. Goldstein, B. J. Schnapp, Bead movement by single kinesin molecules studied with optical tweezers. Nature 348, 348–352 (1990).

7. A. Yildiz, M. Tomishige, R. D. Vale, P. R. Selvin, Kinesin walks hand-over-hand. Science 303, 676–678 (2004).

8. I. A. Telley, P. Bieling, T. Surrey, Obstacles on the microtubule reduce the processivity of Kinesin-1 in a minimal in vitro system and in cell extract. Biophys. J. 96, 3341–3353 (2009).

9. B. Y. Monroy et al., A Combinatorial MAP Code Dictates Polarized Microtubule Transport. Dev. Cell 53, 60–72.e64 (2020).

10. L. S. Ferro et al., Structural and functional insight into regulation of kinesin-1 by microtubule-associated protein MAP7. Science 375, 326–331 (2022).

11. R. Tan et al., Microtubules gate tau condensation to spatially regulate microtubule functions. Nat. Cell Biol. 21, 1078–1085 (2019).

12. V. Siahaan et al., Kinetically distinct phases of tau on microtubules regulate kinesin motors and severing enzymes. Nat. Cell. Biol. 21, 1086–1092 (2019).

13. A. Seitz et al., Single-molecule investigation of the interference between kinesin, tau and MAP2c. EMBO J. 21, 4896–4905 (2002).

14. M. Vershinin, B. C. Carter, D. S. Razafsky, S. J. King, S. P. Gross, Multiple-motor based transport and its regulation by Tau. Proc. Natl. Acad. Sci. U.S.A. 104, 87–92 (2007).

15. R. Dixit, J. L. Ross, Y. E. Goldman, E. L. Holzbaur, Differential regulation of dynein and kinesin motor proteins by tau. Science 319, 1086–1089 (2008).

16. M. D. Weingarten, A. H. Lockwood, S. Y. Hwo, M. W. Kirschner, A protein factor essential for microtubule assembly. Proc. Natl. Acad. Sci. U.S.A. 72, 1858–1862 (1975).

17. G. T. Shubeita et al., Consequences of motor copy number on the intracellular transport of kinesin-1-driven lipid droplets. Cell 135, 1098–1107 (2008).

18. A. G. Hendricks et al., Motor coordination via a tug-of-war mechanism drives bidirectional vesicle transport. Curr. Biol. 20, 697–702 (2010).

19. A. Rai et al., Dynein Clusters into Lipid Microdomains on Phagosomes to Drive Rapid Transport toward Lysosomes. Cell 164, 722–734 (2016).

20. M. Shinitzky, Patterns of lipid changes in membranes of the aged brain. Gerontology 33, 149–154 (1987).

21. R. G. Cutler et al., Involvement of oxidative stress-induced abnormalities in ceramide and cholesterol metabolism in brain aging and Alzheimer’s disease. Proc. Natl. Acad. Sci. U.S.A. 101, 2070–2075 (2004).

22. J. N. Sleigh, A. M. Rossor, A. D. Fellows, A. P. Tosolini, G. Schiavo, Axonal transport and neurological disease. Nat. Rev. Neurol. 15, 691–703 (2019).

23. Q. Li, K. F. Tseng, S. J. King, W. Qiu, J. Xu, A fluid membrane enhances the velocity of cargo transport by small teams of kinesin-1. J. Chem. Phys. 148, 123318 (2018).

24. R. Jiang et al., Microtubule binding kinetics of membrane-bound kinesin-1 predicts high motor copy numbers on intracellular cargo. Proc. Natl. Acad. Sci. U.S.A. 116, 26564–26570 (2019).

25. S. Pyrpassopoulos, H. Shuman, E. M. Ostap, Method for measuring single-molecule adhesion forces and attachment lifetimes of protein-membrane interactions. Methods Mol. Biol. 1046, 389–403 (2013).

26. B. B. McIntosh, E. L. Holzbaur, E. M. Ostap, Control of the initiation and termination of kinesin-1-driven transport by myosin-Ic and nonmuscle tropomyosin. Curr. Biol. 25, 523–529 (2015).

27. R. Grover et al., Transport efficiency of membrane-anchored kinesin-1 motors depends on motor density and diffusivity. Proc. Natl. Acad. Sci. U.S.A. 113, E7185–E7193 (2016).

28. C. Herold, C. Leduc, R. Stock, S. Diez, P. Schwille, Long-range transport of giant vesicles along microtubule networks. Chemphyschem 13, 1001–1006 (2012).

29. S. R. Nelson, K. M. Trybus, D. M. Warshaw, Motor coupling through lipid membranes enhances transport velocities for ensembles of myosin Va. Proc. Natl. Acad. Sci. U.S.A. 111, E3986–E3995 (2014).

30. K. Svoboda, S. M. Block, Force and velocity measured for single kinesin molecules. Cell 77, 773–784 (1994).

31. J. Xu et al., Casein kinase 2 reverses tail-independent inactivation of kinesin-1. Nat. Commun. 3, 754 (2012).

32. S. Klumpp, R. Lipowsky, Cooperative cargo transport by several molecular motors. Proc. Natl. Acad. Sci. U.S.A. 102, 17284–17289 (2005).

33. J. Xu, Z. Shu, S. J. King, S. P. Gross, Tuning multiple motor travel via single motor velocity. Traffic 13, 1198–1205 (2012).

34. Q. Li, S. J. King, A. Gopinathan, J. Xu, Quantitative Determination of the Probability of Multiple-Motor Transport in Bead-Based Assays. Biophys. J. 110, 2720–2728 (2016).

35. E. M. Jones et al., Interaction of tau protein with model lipid membranes induces tau structural compaction and membrane disruption. Biochemistry 51, 2539–2550 (2012).

36. A. Parker, K. Miles, K. H. Cheng, J. Huang, Lateral distribution of cholesterol in dioleoylphosphatidylcholine lipid bilayers: cholesterol-phospholipid interactions at high cholesterol limit. Biophys. J. 86, 1532–1544 (2004).

37. A. R. Rogers, J. W. Driver, P. E. Constantinou, D. Kenneth Jamison, M. R. Diehl, Negative interference dominates collective transport of kinesin motors in the absence of load. Phys. Chem. Chem. Phys. 11, 4882–4889 (2009).

38. N. D. Derr et al., Tug-of-war in motor protein ensembles revealed with a programmable DNA origami scaffold. Science 338, 662–665 (2012).

39. K. Furuta et al., Measuring collective transport by defined numbers of processive and nonprocessive kinesin motors. Proc. Natl. Acad. Sci. U.S.A. 110, 501–506 (2013).

40. S. R. Norris et al., A method for multiprotein assembly in cells reveals independent action of kinesins in complex. J. Cell Biol. 207, 393–406 (2014).

41. M. H. Hinrichs et al., Tau protein diffuses along the microtubule lattice. J. Biol. Chem. 287, 38559–38568 (2012).

42. D. P. McVicker, G. J. Hoeprich, A. R. Thompson, C. L. Berger, Tau interconverts between diffusive and stable populations on the microtubule surface in an isoform and lattice specific manner. Cytoskeleton 71, 184–194 (2014).

43. J. D. Knight, M. G. Lerner, J. G. Marcano-Velazquez, R. W. Pastor, J. J. Falke, Single molecule diffusion of membrane-bound proteins: window into lipid contacts and bilayer dynamics. Biophys. J. 99, 2879–2887 (2010).

44. F. A. Thomas, I. Visco, Z. Petrasek, F. Heinemann, P. Schwille, Diffusion coefficients and dissociation constants of enhanced green fluorescent protein binding to free standing membranes. Data Brief 5, 537–541 (2015).

45. C. Leduc et al., Cooperative extraction of membrane nanotubes by molecular motors. Proc. Natl. Acad. Sci. U.S.A. 101, 17096–17101 (2004).

46. L. Amos, A. Klug, Arrangement of subunits in flagellar microtubules. J. Cell Sci. 14, 523–549 (1974).

47. M. J. Schnitzer, S. M. Block, Kinesin hydrolyses one ATP per 8-nm step. Nature 388, 386–390 (1997).

48. E. H. Kellogg et al., Near-atomic model of microtubule-tau interactions. Science 360, 1242–1246 (2018).

49. D. P. McVicker, L. R. Chrin, C. L. Berger, The nucleotide-binding state of microtubules modulates kinesin processivity and the ability of Tau to inhibit kinesin-mediated transport. J. Biol. Chem. 286, 42873–42880 (2011).

50. J. L. Stern, D. V. Lessard, G. J. Hoeprich, G. A. Morfini, C. L. Berger, Phosphoregulation of Tau modulates inhibition of kinesin-1 motility. Mol. Biol. Cell 28, 1079–1087 (2017).

51. B. Y. Monroy et al., Competition between microtubule-associated proteins directs motor transport. Nat. Commun. 9, 1487 (2018).

52. C. Lebrand et al., Late endosome motility depends on lipids via the small GTPase Rab7. EMBO J. 21, 1289–1300 (2002).

53. E. B. Neufeld et al., The ABCA1 transporter modulates late endocytic trafficking: insights from the correction of the genetic defect in Tangier disease. J. Biol. Chem. 279, 15571–15578 (2004).

54. D. R. Klopfenstein, M. Tomishige, N. Stuurman, R. D. Vale, Role of phosphatidylinositol(4,5)bisphosphate organization in membrane transport by the Unc104 kinesin motor. Cell 109, 347–358 (2002).

55. S. E. Encalada, L. Szpankowski, C. H. Xia, L. S. Goldstein, Stable kinesin and dynein assemblies drive the axonal transport of mammalian prion protein vesicles. Cell 144, 551–565 (2011).

56. J. Kerssemakers, J. Howard, H. Hess, S. Diez, The distance that kinesin-1 holds its cargo from the microtubule surface measured by fluorescence interference contrast microscopy. Proc. Natl. Acad. Sci. U.S.A. 103, 15812–15817 (2006).

57. G. Woehlke et al., Microtubule interaction site of the kinesin motor. Cell 90, 207–216 (1997).

58. G. Skiniotis et al., Modulation of kinesin binding by the C-termini of tubulin. EMBO J. 23, 989–999 (2004).

59. B. J. Grant et al., Electrostatically biased binding of kinesin to microtubules. PLoS Biol. 9, e1001207 (2011).

60. W. H. Liang et al., Microtubule Defects Influence Kinesin-Based Transport In Vitro. Biophys. J. 110, 2229–2240 (2016).

61. E. P. Karasmanis et al., Polarity of Neuronal Membrane Traffic Requires Sorting of Kinesin Motor Cargo during Entry into Dendrites by a Microtubule-Associated Septin. Dev. Cell 46, 518–524 (2018).

62. M. W. Gramlich et al., Single Molecule Investigation of Kinesin-1 Motility Using Engineered Microtubule Defects. Sci. Rep. 7, 44290 (2017).

63. M. Metivier et al., Dual control of Kinesin-1 recruitment to microtubules by Ensconsin in Drosophila neuroblasts and oocytes. Development 146, dev171579 (2019).

64. P. J. Hooikaas et al., MAP7 family proteins regulate kinesin-1 recruitment and activation. J. Cell Biol. 218, 1298–1318 (2019).

65. J. C. Larcher, D. Boucher, S. Lazereg, F. Gros, P. Denoulet, Interaction of kinesin motor domains with alpha-and beta-tubulin subunits at a tau-independent binding site. Regulation by polyglutamylation. J. Biol. Chem. 271, 22117–22124 (1996).

66. G. Liao, G. G. Gundersen, Kinesin is a candidate for cross-bridging microtubules and intermediate filaments. Selective binding of kinesin to detyrosinated tubulin and vimentin. J. Biol. Chem. 273, 9797–9803 (1998).

67. Y. Konishi, M. Setou, Tubulin tyrosination navigates the kinesin-1 motor domain to axons. Nat. Neurosci. 12, 559–567 (2009).

68. N. Kaul, V. Soppina, K. J. Verhey, Effects of alpha-tubulin K40 acetylation and detyrosination on kinesin-1 motility in a purified system. Biophys. J. 106, 2636–2643 (2014).

69. T. E. Smith et al., Single-molecule inhibition of human kinesin by adociasulfate-13 and -14 from the sponge Cladocroce aculeata. Proc. Natl. Acad. Sci. U.S.A. 110, 18880–18885 (2013).

70. B. C. Carter, G. T. Shubeita, S. P. Gross, Tracking single particles: a user-friendly quantitative evaluation. Phys. Biol. 2, 60–72 (2005).

## SI References

1. S. M. Block, L. S. Goldstein, B. J. Schnapp, Bead movement by single kinesin molecules studied with optical tweezers. Nature 348, 348–352 (1990).

2. K. Svoboda, S. M. Block, Force and velocity measured for single kinesin molecules. Cell 77, 773–784 (1994).

3. E. M. Jones et al., Interaction of tau protein with model lipid membranes induces tau structural compaction and membrane disruption. Biochemistry 51, 2539–2550 (2012).

